# Native, full-length, refolding-assisted purification of TDP-43 compatible with BSL-2 safety regulations

**DOI:** 10.64898/2025.12.10.693404

**Authors:** Subhasis Dehury, Satyam Tiwari, Paolo De Los Rios

**Affiliations:** Institute of Physics, School of Basic Sciences, École Polytechnique Fédérale de Lausanne - EPFL, Lausanne, Switzerland; Institute of Bioengineering, School of Life Sciences, École Polytechnique Fédérale de Lausanne - EPFL, Lausanne, Switzerland

## Abstract

TAR DNA-binding protein 43 (TDP-43) is a prion-like RNA-binding protein that plays a key role in amyotrophic lateral sclerosis and frontotemporal dementia. Producing full-length TDP-43 consistently is thus relevant for its in vitro studies and yet it remains challenging, especially with the current requirement to work under biosafety level-2 (BSL-2) containment due to new safety regulations for Prion-like and amyloidogenic proteins. Here we describe a refolding-assisted purification protocol for TDP-43 from soluble fraction that can be implemented with basic equipment in standard BSL-2 laboratories. Expression in Escherichia coli is followed by IMAC-capture on an EDTA/DTT-tolerant Ni-NTA resin under 4 M urea, then on-column refolding via a gradient urea wash using resin-limiting conditions that favour the binding to high-affinity His-tagged protein. After SUMO solubility tag removal, the preparation is monitored by a robust quality-control pipeline: SDS-PAGE and immunoblotting for integrity and purity, mass photometry for oligomeric state, far-UV circular dichroism for secondary structure, fluorescence anisotropy for native functional assays, and light-scattering for stability and aggregation propensity measurements. A concise BSL-2 standard operating procedure specifies containment, decontamination, and waste handling for prion-like proteins. This protocol enables safe, cost-effective, and reproducible access to native-like full-length TDP-43 and is readily adaptable to other prion-like aggregation-prone proteins.

**Highlights:**

- Presents a single-step, gravity-based Immobilized metal affinity (IMAC) purification with an optimized on-column refolding strategy.
- Includes a detailed BSL-2 Standard Operating Procedure (SOP) with specific decontamination contact times and waste management protocols.
- Validation of protein quality, oligomeric state, and functional characterisation

## 1. Introduction

Prions are aberrantly folded proteins that can be toxic to cells, disrupting function and promoting cell death[1]. They are considered infectious because their pathogenic conformation spreads within and between organisms by seeded, self-templated conversion of normally folded proteins into the same misfolded and aggregated structure[2,3]. In mammals, prions are responsible for a group infectious neurodegenerative diseases, also called Transmissible Spongiform Encephalopathies (TSEs)[4]. A protein qualifies as a prion in the strict sense when it satisfies three main criteria: infectivity as a protein-only agent, such that purified, nucleic-acid-free aggregates transmit disease and can propagate to a new host; self-templated propagation, whereby the misfolded conformer converts the native protein to the same misfolded state; and host-to-host transmissibility associated with pathology under experimental or natural conditions, as in mammalian TSEs[5,6].

Importantly the ability to template misfolding can occur without established host-to-host infectivity. Proteins that display this behaviour are usually termed prion-like or proteopathic seeds[2,7]. TDP-43 (Transactivation response (TAR), DNA-binding protein of 43 kDa) was first isolated as a transcriptional inactivator binding to the TAR DNA element of the HIV-1 virus [8](Fig. 1a), Its ubiquitinated and phosphorylated, forms plays a role in ALS (Amyotrophic lateral sclerosis) and FTD (Frontotemporal dementia) [9]. TDP-43 is composed of 414 amino acids, it has an N-terminus domain (NTD) of unknown function, two functional RNA-binding domains (RRM1 and RRM2) and a prion-like domain (PrLD) responsible for protein-protein interactions (Fig. 1a, d)[10]. TDP-43 falls under the category of prion-like protein because of its PrLD (Prion-like domain) domain in the C-terminus [11]. TDP-43 exhibits templated-seeded aggregation and cell-to-cell spread in cell culture and animal models, and this behaviour has been linked to its pathology in explaining staged ALS/FTD progression [12,13]. It has not been shown that TDP-43 infects naturally or iatrogenically between humans or hosts through protein alone (no TSE-like transmission established)[14], Therefore, it is currently classified only as a prion-like protein. Studying the behaviour of TDP-43 in vitro is therefore essential for understanding the molecular mechanisms of these diseases and for developing potential therapeutics. Such studies are critically dependent on the availability of high-quality, pure, and functionally active proteins, whose production must comply with the new safety standards required for prion-like proteins, such as biosafety level-2( BSL) containment[15].

**Fig. 1.**
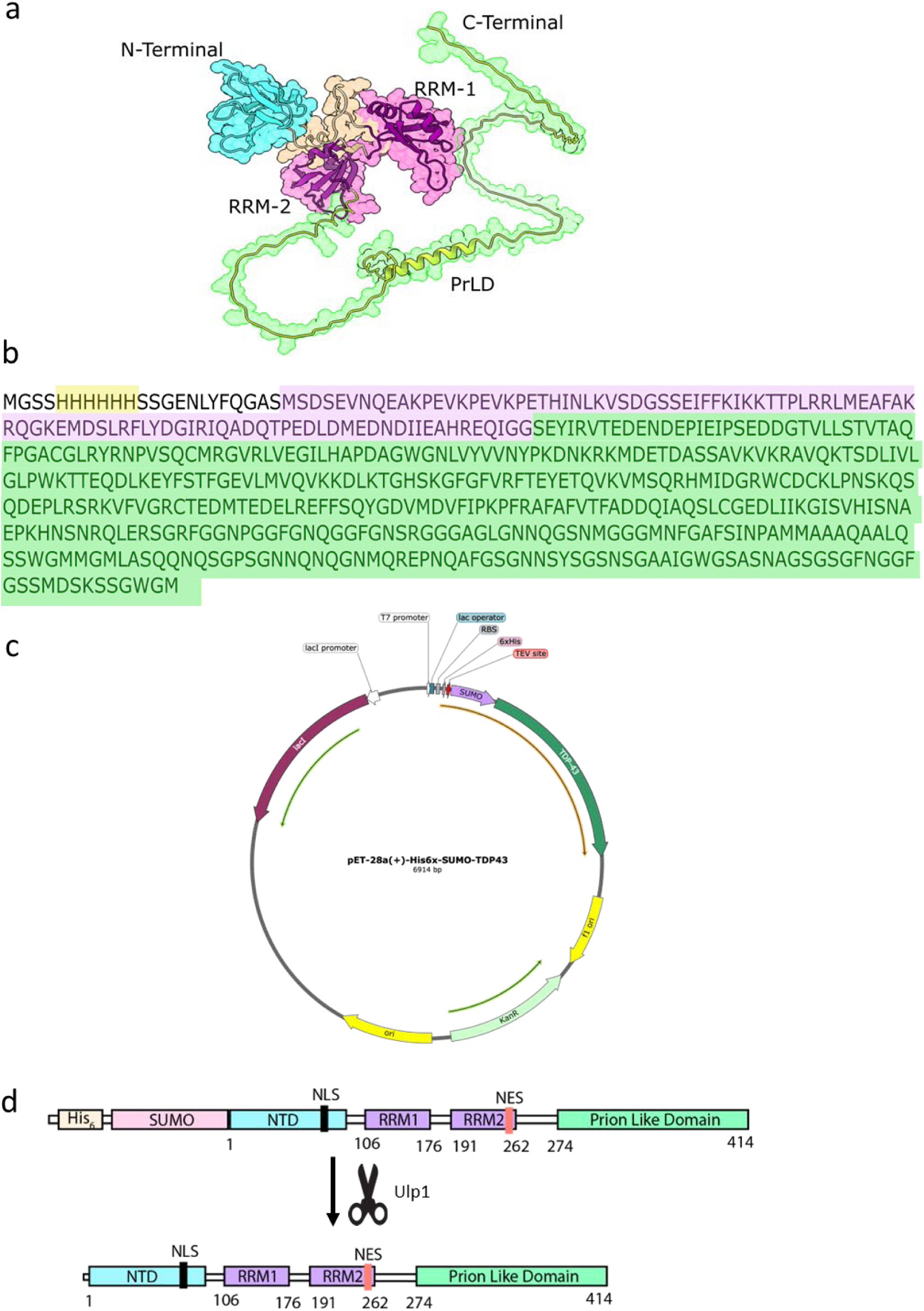
Domain organization, amino acid sequence, and vector map of TDP-43: a) AlphaFold structure of TDP-43: The structure, rendered as a space-filling model in Chimera. b) Amino acid sequence of His₆-SUMO-TDP-43: The full amino acid sequence is color-coded by component: His₆ tag (yellow), SUMO tag (pink), and full-length TDP-43 (green). c) Final Vector Map: plasmid map of the His₆-SUMO-TDP-43 construct in the pET-28a(+) backbone. d) Domain Organization of TDP-43: Conserved domains identified via uniprot. NTD (N-terminal domain) is shown in cyan, RRM (RNA recognition motifs) in magenta, and PrLD (Prion-like domain) in lemon green, NLS (Nuclear localizations signal) and NES (Nuclear export signal) in black and pink respectively.

In this work, we show end-to-end native purification of full-length TDP-43. The main challenge of TDP-43 purification is its prion-like nature, which requires a BSL-2 lab with new biosafety regulations and often a designated FPLC (Fast performance liquid chromatography) system. Even with a designated FPLC, vigorous cleaning is required due to the protein’s high tendency to aggregate quickly. Here, we demonstrate that, by using an optimized protocol, we can purify TDP-43 with a single-step, gravity-based immobilised metal affinity chromatography (IMAC) method, embedding safety controls aligned with guidance on handling prion-like proteins, including risk assessment, containment, decontamination, and waste management.

## 2. Description of Methods and Results

### 2.1 Cloning strategy and expression Plasmid vector Construct

A His₆-SUMO-TDP43 construct (Fig. 1b,c) was cloned into the pET28(a) expression vector (GeneScript). The His₆-SUMO-TDP43 plasmid was verified by full-length plasmid sequencing. We chose the N-terminal His₆-SUMO (Small Ubiquitin-like Modifier), ∼17 kDa, tag for its known solubility-enhancing properties (since TDP-43 is highly aggregation-prone). Its cleavage by the Ulp-1 (ubiquitin-like proteases) protease is highly specific [16]. This allows precise removal of the entire tag, leaving an intact, native N-terminus on the target protein (Fig. 1d). It has significant advantage over other large tags like MBP (∼42 kDa) or GST (∼26 kDa), which often leave extra amino acids. In-house Ulp-1 production significantly reduces commercial costs.

### 2.2 Protein expression and purification procedure

For the expression of His₆-SUMO-TDP43, the plasmid was transformed into the *E. coli* strain BL21-CodonPlus (DE3)-RIPL using kanamycin (50 µg/ml) and chloramphenicol (35 µg/ml). An overnight starter culture (20ml) from a single colony, grown at 25°C, was used to inoculate 2 liters of LB medium, which was further grown at 25°C to an OD₆₀₀ of 0.3-0.4. The culture was then cooled to 16°C for 60 minutes without shaking (a crucial step), and protein expression was induced with 0.2 mM IPTG overnight at 16°C (∼16 hours). Whole-cell lysates show induction-dependent appearance of the target band (UI vs I), with the protein partitioning largely into the soluble fraction (Sup) and moderate loss in the flow-through (FT) (Fig. 3a).

All subsequent steps were conducted at 4°C unless otherwise specified (critical step). Cells were harvested by centrifugation at 5’000 g for 15 min (pre-cooled rotor-JLA-9.1000 at 4°C), and the pellet was washed with chilled buffer (30 mM HEPES-KOH, pH 7.5, 150 mM KCl, 5% Glycerol). The cell pellet was resuspended in 80 mL of lysis buffer (50 mM HEPES-KOH pH 7.5, 500 mM NaCl, 5 mM EDTA, 10 mM Imidazole, 2 mM PMSF, 2 tablets cOmplete™ Protease Inhibitor Cocktail, 2 mM DTT, 40 mg lysozyme , 200 µg DNase I , 200 µg RNase , 10% glycerol). Cell lysis was performed at 18’000 psi using microfluidizer, and the lysate was clarified by ultracentrifugation at 25’000 g for 30 min (pre-cooled rotor Type 45 Ti at 4°C, Beckman Coulter Optima XE-90 Ultracentrifuge).

Following clarification, the supernatant containing the soluble TDP-43 was securely transferred to BSL-2 laboratory (Fig. 2 schematic).

**Fig 2.**
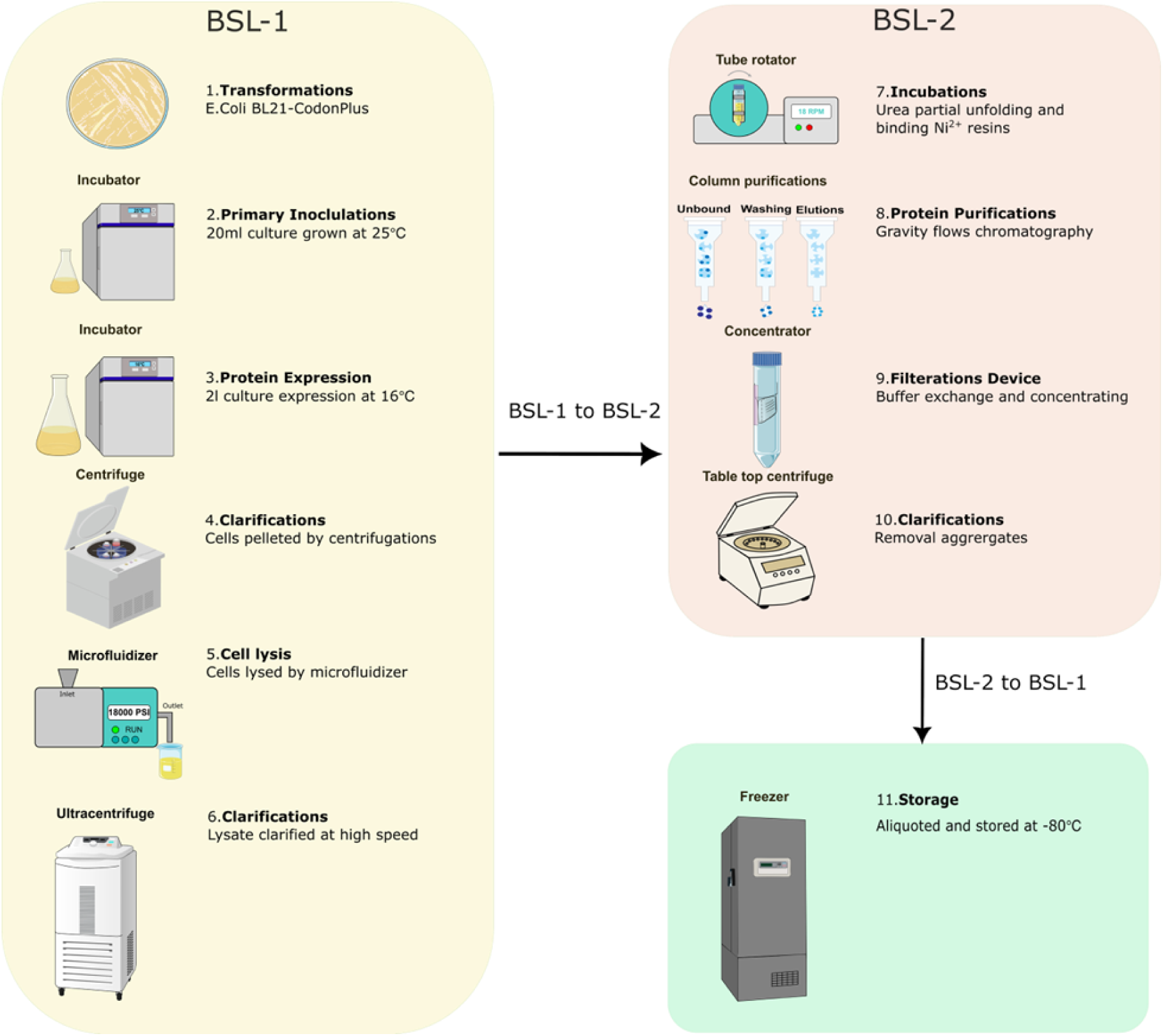
Schematic of the His₆-SUMO-TDP43 purification workflow. The diagram outlines the major steps, clearly delineating procedures performed under BSL-1 containment (yellow box) and those requiring the BSL-2 environment (pink box). The figure was created using Inkscape with icons from the Biocons plugin.

**Fig. 3.**
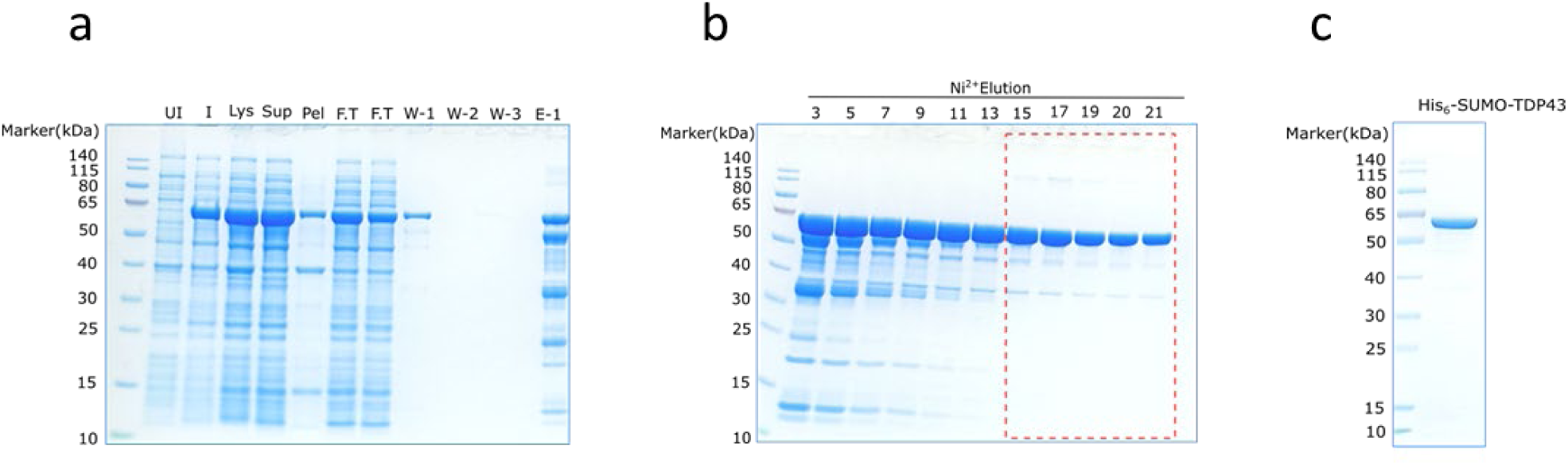
Expression and IMAC-Purification of His_6_-SUMO-TDP43. Coomassie-stained 12% SDS-PAGE gel analysing various fractions from the Ni-NTA gravity-flow purification. Lanes are designated as follows: M, Molecular weight marker (kDa); UI-Uninduced; I-Induced; Lys-Lysate; Sup-Supernatant; Pel-pellet, FT-Flow-through; W-Wash; E1-E4, Elution fractions. For each lane, ∼6 µg total protein was loaded. The target protein is successfully purified, indicated by the prominent band at the expected molecular weight of ∼58 kDa in the elution lanes. c. Pooled-fraction purity.

The supernatant was filtered using a 0.45µm PVDF filter. To prevent aggregation and enhance binding of His_6_-SUMO-TDP43 to the Ni^2+^-affinity resin, 4 M urea was added to the supernatant, and the solution was incubated on a tube revolver rotator for 30 minutes at 4°C. The solution was filtered with 0.2 µm PES filter and incubated with 1.5 ml of pre-equilibrated EDTA-compatible Ni²⁺-NTA beads for 2 hours on a tube revolver rotator. Notably, a smaller volume of beads (1.5 ml) was used relative to the estimated protein yield; this was intentionally done to increase binding competition and select for the highest-affinity His_6_-tagged protein only. It is also crucial to use these specific EDTA-compatible Ni^2+^-Beads from Roche, as our lysis and wash buffers contain EDTA and DTT, which can inactivate standard Ni²⁺-beads.

After binding, the Ni^2+^-bead slurry was transferred to an empty PD-10 gravity-flow column. Unbound (flowthrough) was collected. On-column refolding was carried out by washing the TDP-43 bound Ni²⁺ resin with 50 ml of base wash buffer (30 mM HEPES-KOH pH 7.5, 250 mM KCl, 20 mM imidazole, 2 mM DTT, 1 mM PMSF, 2 mM EDTA, 1× protease inhibitor tablet, 5% glycerol) applied sequentially with decreasing urea concentrations: first 2 M, then 1 M, and finally 0 M urea (50 ml each was buffer with different urea concentration). This stepwise removal of denaturant promotes gradual refolding of the bound His₆-SUMO-TDP43 while it remains on the column.

The protein was eluted with elution buffer (30 mM HEPES-KOH pH 7.5, 200 mM KCl, 300 mM Imidazole, 2 mM DTT, 1 mM PMSF, 2 mM EDTA, 1x protease inhibitor tablet, 5% glycerol) into 2ml protein low-binding tubes, 2ml each fraction, until all protein was eluted completely by quickly checking the colour change of Bradford reagent. (10ul eluted sample + 90ul Bradford reagent in a 96-well, transparent plate). Elution with imidazole yielded a preparation enriched in a single dominant ∼58 kDa species corresponding to His₆-SUMO-TDP43 (Fig. 3c). Pure fractions (fraction 15 to 21, marked in red Fig. 3b) were pooled, buffer-exchanged, and concentrated using 100 kDa concentrator to ∼20 µM with final storage buffer (30 mM HEPES-KOH pH 7.5, 100 mM KCl, 2 mM DTT, 5% glycerol).

Notably, using a 100 kDa MWCO filter for the ∼58 kDa protein resulted in minimal protein loss and appeared to reduce aggregation during concentration process. The concentrated samples were centrifuged again (10,000 g, 10 min, 4°C) to ensure no big oligomers/aggregates before aliquoting in 1.5ml Protein LoBind tube and stored at −80°C. Densitometry across the pooled eluate gives ∼90% purity and a stepwise yield of 4.6 mg per litre culture . These meet the release criteria for downstream tag removal (purity ∼90%, pooled yield ∼**10** mg) (Fig 3c).

All procedures within the BSL-2 facility were performed with strict adherence to safety protocols. Samples were moved between labs using double container protection (secondary containment). Any aggregated samples were discarded in designated BSL-2 trash bins (30 Litre blue boxes certified UN 3291, OMoD code 18 01 03), since this is special waste, it has to comply with the Federal Act on the Protection of the Environment (EPA, ref. 814.01) and the Ordinance on the Transport of Waste (OMoD, ref. 814.610), while also adhering to the biosafety standards outlined in the NIH Guidelines for Research Involving Recombinant or Synthetic Nucleic Acid Molecules and the BMBL(Biosafety in Microbiological and Biomedical Laboratories) (6th Edition). Spills on solid surfaces were decontaminated with 1% Hellmanex III and 2% SDS solutions, and all liquid waste was inactivated with 1 M NaOH before disposal.

### 2.3 Sumo tag removal and truncation analysis

To test SUMO-tag cleavage and confirm product identity, a time-course experiment was performed. His₆-SUMO-TDP43 at 5 µM concentration was incubated with 1 µM Ulp1 protease at room temperature, with samples taken at various time points and run in SDS-PAGE electrophoresis. SUMO removal was efficient, with less than 5 minutes required for 100% cleavage efficiency, and it preserved full-length TDP-43. Coomassie 12% SDS–PAGE shows loss of the ∼58 kDa precursor and appearance of two products: TDP-43 (∼43 kDa) and the released His₆-SUMO ∼17 kDa, with Ulp1 at ∼27 kDa (Fig. 4a). To further test truncation, SUMO-cleaved sample was taken for western blot analysis. The sample was run on a 12% SDS-PAGE gel and transferred onto a nitrocellulose membrane (0.4 µm), at 90V for 90 minutes using 1x Tris-Glycine buffer in a cold room. The membrane was blocked for 1 hour in 5% BSA in TBST, then incubated with anti-TDP43 polyclonal rabbit antibody (1:5000, antibody recognizes the full length TDP-43) overnight at 4°C. The membrane was washed, three times, 5 min each, with PBST buffer and incubated with an anti-rabbit HRP-conjugated secondary antibody (1:10,000) for 1 hour at room temperature. The blot was developed using the ECL western blotting analysis system and visualized on an iBrightFL1000. Immunoblotting with anti-TDP-43 shows a single band around 44 kDa (Fig. 4b), confirming the size of full-length TDP-43 with no detectable truncation of purified protein.

**Fig. 4.**
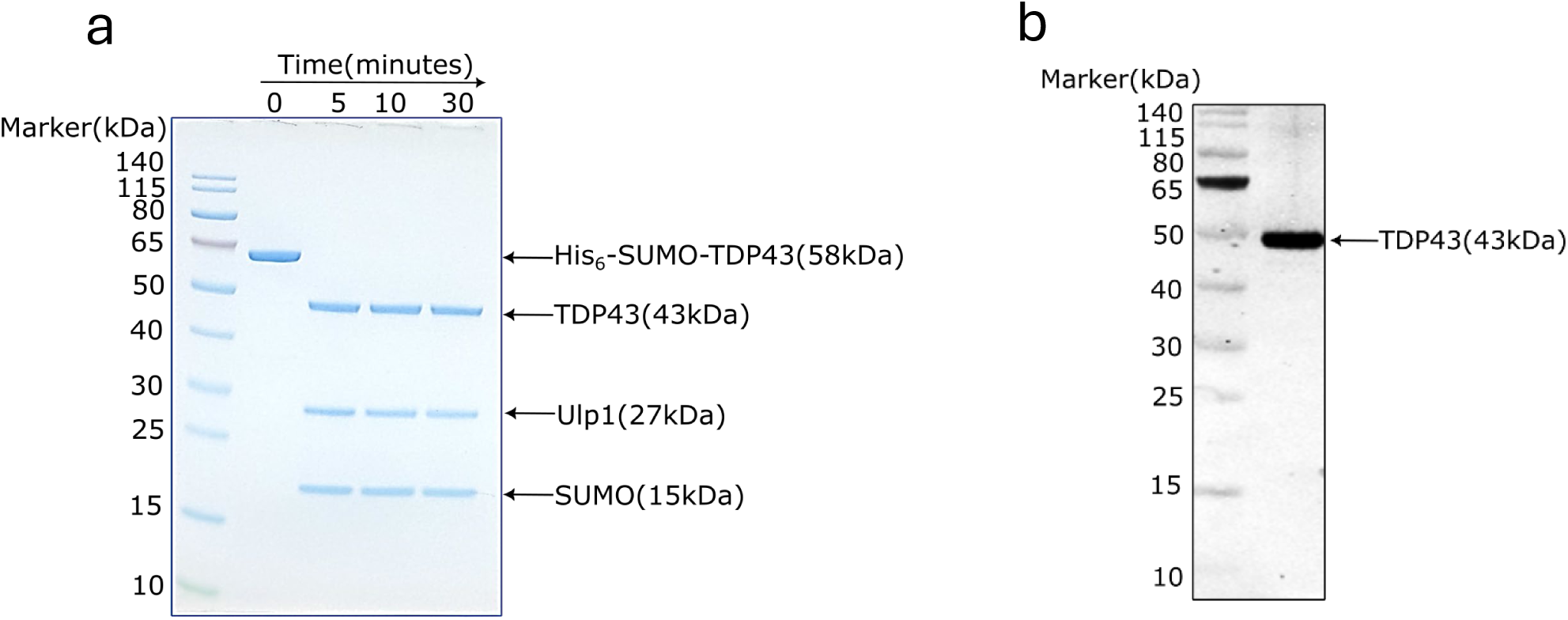
Time-course analysis of SUMO-tag cleavage from TDP-43. (a) A time-course cleavage experiment was performed by incubating 5 µM His_6_-SUMO-TDP43 with 1 µM Ulp1 protease at RT. Samples were taken at different time points (5, 10, and 30 minutes) and analyzed on a 12% SDS-PAGE gel to monitor the reaction. Bands corresponding to TDP-43 (∼43 kDa), Ulp1 (∼27 kDa), and His₆–SUMO (∼17 kDa) are indicated. 5 µg total protein loaded per lane, (b) A western blot was performed on a cleaved sample using a polyclonal anti-TDP43 antibody to confirm that no non-specific cleavage of TDP-43 occurred during the removal of the SUMO tag by Ulp1.

### 2.4 Mass photometry reports predominantly monomeric TDP-43 before and after SUMO tag removal

Mass photometry was used to determine the oligomeric state of the purified TDP-43 providing single-particle level resolution using a TwoMP instrument from Refeyn. The TDP-43 sample was first thawed within a BSL-2 facility and then securely transported in a double-protected container to the BSL-1 lab to get access the Refeyn mass photometer. This analysis is crucial given TDP-43’s tendency to form various oligomeric species. Proteins were diluted to 50 nM in 50 mM Tris-HCl, pH 7.5, 100 mM NaCl, 2 mM DTT before measurements. Bovine serum albumin (BSA) was used for mass calibration (Fig. 5a,b). Single-particle mass photometry confirmed that both the His₆-SUMO-TDP43 precursor and the tag-free product are largely monomeric in storage buffer: His₆-SUMO–TDP43 was detected as a single major population centered at ∼58 kDa, accounting for ∼ 94% of 1465 events (Fig. 5c), and after Ulp1 cleavage the mass distribution shifts to ∼48-50 kDa with monomer (∼82% of 822 events) (Fig. 5d). Minor higher-mass events account for the balance and represent less than the 20% acceptance limit. After the measurement, the used glass slides were carefully collected and disposed of in a designated BSL-2 waste container.

**Fig. 5.**
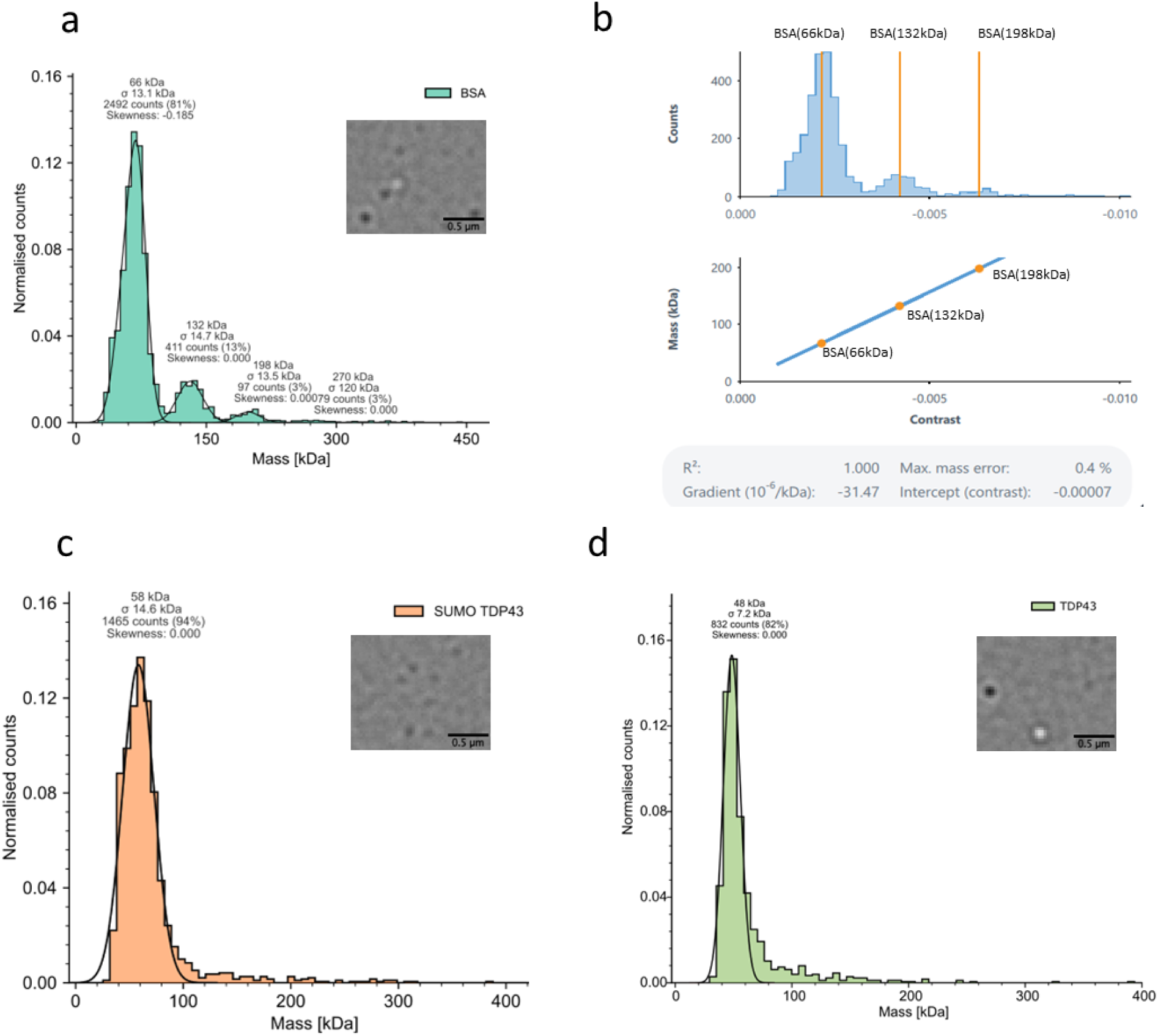
Mass Photometry analysis of the TDP43 oligomeric state. Single-particle mass photometry (iSCAT) was performed using a TwoMP mass photometer to determine the native oligomeric state and mass of the purified protein. BSA was used as the calibration standard. Movies were recorded for 60 seconds, and the resulting mass histograms were fitted with Gaussian functions. Samples were replicated three times a) Mass distribution of the BSA standard, showing its characteristic monomeric 66 kDa, dimeric132 kDa, trimeric 198 kDa peak and its oligomeric species inset, representative landing image. b) The linear calibration curve generated by plotting the known BSA masses in DiscoverMP software. slope ∼1.00, intercept ∼0, max mass error ∼0.4% c) Purified 6xHis-SUMO-TDP43 reveals a single major population with a measured mass of approximately 58 kDa, consistent with a dominant monomeric species. inset, representative landing image. d) Mass distribution of TDP43 following enzymatic removal of the SUMO domain shows a predominant species at approximately 48 kDa, corresponding to the expected molecular weight of the monomeric, tag-free protein. inset, representative landing image. Each samples were analyzed on microscope coverslips with silicone cassettes in a buffer containing 50 mM Tris pH 7.5, 100 mM NaCl, and 2 mM DTT, which had been passed through a 0.2 µm syringe filter. Each sample was recorded for 60 second per movie, n = 3 independent preparations. Gaussian fits were applied to event histograms

### 2.5 Far-UV CD shows a folded, mixed α/β spectrum for SUMO–TDP-43

To control if the secondary structure of the protein is preserved, Far-UV CD spectra were recorded on a Chirascan V100 instrument. TDP-43 (20 µM stock concentration) was diluted 1:10 in Milli-Q water to a final concentration of 2 µM. A 250 µl sample was loaded into a 1 mm path-length quartz cuvette. The resulting CD spectrum shows the canonical α/β signature (Fig. 6). Data displays negative minima around 208 nm and 224 nm, indicative of α-helical content, in agreement with previous work [17,18], and a negative shoulder around 216 nm, indicative of β-sheet content, confirming a folded, mixed-structure protein. As a safety precaution, the cuvette was sealed with parafilm to prevent any accidental spillage of the BSL-2 sample.

**Fig. 6.**
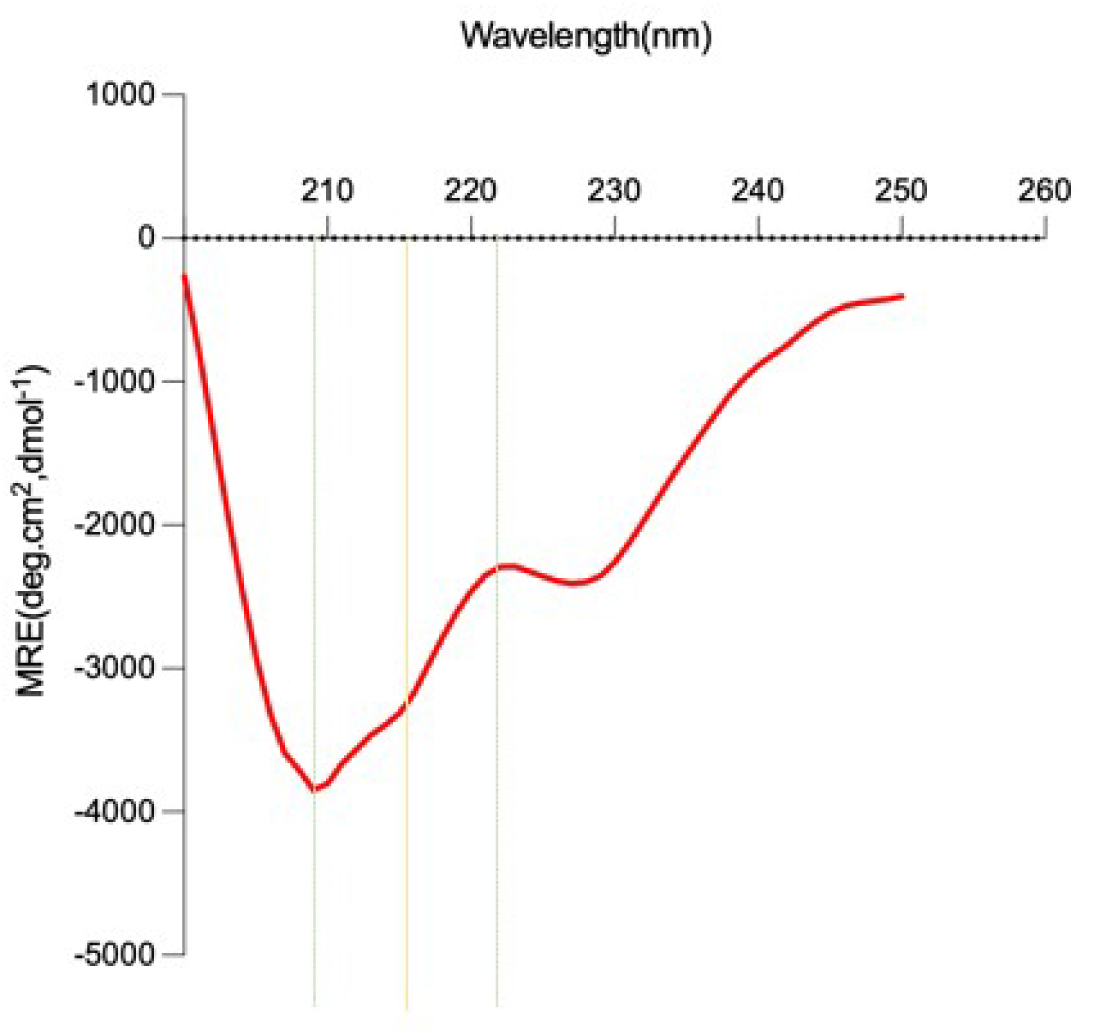
Secondary structure analysis by Circular Dichroism. The CD spectrum was recorded on a Chirascan V100 instrument. SUMO-TDP43 (1 µM) was scanned from 250 nm to 200 nm at 23°C (1 nm bandwidth, 1 nm step size), with a 30-second scan time. Three scans were collected per sample and averaged; a corresponding buffer blank was recorded and subtracted from the average sample spectrum. The resulting spectrum, displays characteristic negative minima around 208 nm and 224 nm (indicative of α-helical content) and a negative shoulder around 216 nm (indicative of β-sheet content), confirming a folded, mixed-structure protein. The final data was converted to Mean Residue Ellipticity (MRE) deg·cm²·dmol⁻¹ using the protein concentrations and number of amino acid residues in chirascan software.

### 2.6 Purified TDP-43 is functionally active and binds specifically to TG-rich ssDNA

To validate that the purified protein was in a native, functionally active state, its ability to bind a known DNA partner was assayed [18,19]. TDP-43 is known to bind TG-rich sequences. FAM-linked ssDNA (TG)_4_ and non-specific (AC)_4_ (as a control) [19,20] was used. FAM-ssDNA (50nM) dissolved in nuclease free water and titrated with increasing amount (0-10 µM) of His₆-SUMO-TDP43 into a 384-well black clear-bottom plate in 50 mM Tris-HCl pH 7.5, 100 mM NaCl, 2 mM DTT buffer. The plate was sealed with optical adhesive covers, incubated for 15 minutes at 25°C, and transported in a double container to the Tecan Spark plate reader for measurements. Measurement done by Excitation at 480 nm (bandwidth: 20 nm); Emission at 525 nm (bandwidth: 20 nm). The anisotropy calculations done using parallel and perpendicular fluorescence intensity values given by the following equations:

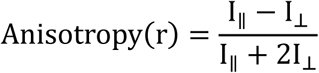

Where I_∥_ equals the emission intensity of the polarized light parallel to the plane of excitation and I_⟂_ equals the emission intensity of the polarized light perpendicular to the plane of excitation.

Consistent with previous reports[18], TDP-43 preferentially binds (TG)-rich sequences. Our purified His₆-SUMO-TDP43 showed a clear, dose-dependent increase in anisotropy when titrated against a (TG)₄ ssDNA sequence, yielding a dissociation constant (K_d_) of **∼**1.8 µM. Conversely, no significant binding was observed when the protein was titrated against a non-specific (AC)₄ ssDNA control sequence (Fig. 7a). These results demonstrate that the purified protein is functionally active and retains its specific binding preference.

**Fig. 7.**
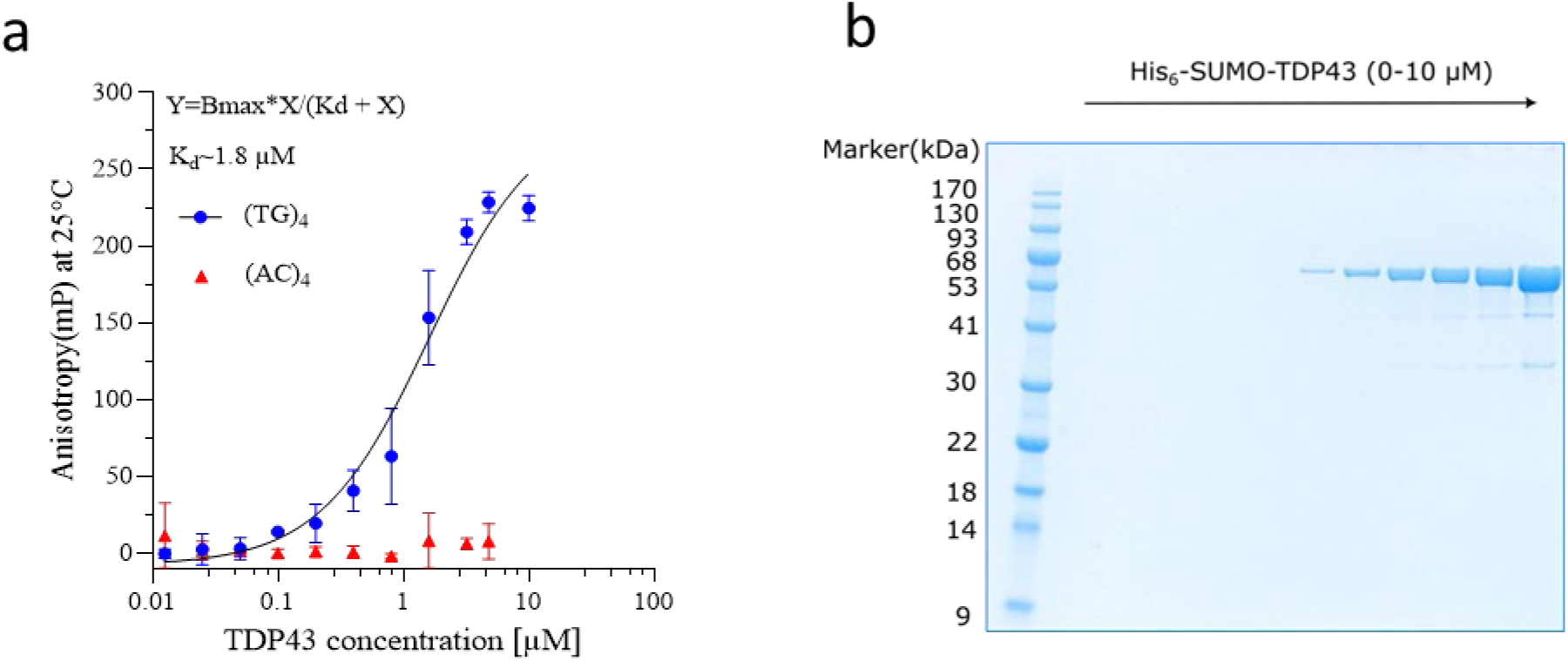
Determination of functional interaction by DNA binding using Fluorescence Anisotropy. a) Dose response of His₆-SUMO-TDP-43 (0-10 µM) to 5′-FAM-labeled ssDNA (50 nM) at 25 °C. TG-rich ssDNA (TG)_4_, a known binder and a non-specific control ssDNA (AC)_4_. Data points represent the mean of three repeats, and the solid line shows a one-site specific binding in a nonlinear equation fit using GraphPad Prism. Measurements were performed using excitation at 480 nm (20 nm bandwidth) and emission at 525 nm (20 nm bandwidth) a gain value of 100, dichroic 510 mirror, and a G-value of 1, at 25°C in TECAN Spark. b) Representative 12% SDS-PAGE of the reaction mixtures used in (a), confirming constant protein integrity across the titration series (0-10 µM).

### 2.7 SUMO removal triggers rapid aggregation detected by Light scattering analysis

The stability and aggregation propensity of TDP-43, a hallmark of its pathology, was initiated by cleaving the solubility-enhancing SUMO tag with Ulp1 protease. 5 µM of His₆-SUMO-TDP43 was incubated with the protease in a 384-well black clear-bottom plate in 50 mM Tris-HCl pH 7.5, 100 mM NaCl, 2 mM DTT at 37 °C. The plate was sealed with optical adhesive film. Light scattering was measured for 120 minutes at 37°C in a Tecan Spark plate reader at 360 nm wavelength. Ulp1 addition to His₆-SUMO-TDP43 causes a sharp rise in turbidity at 360 nm, consistent with aggregation of tag-free TDP-43 at higher temperature. (Fig. 8a, blue). The SUMO-tagged protein without Ulp1 shows only a slow, shallow increase (red), and Ulp1 alone remains near baseline (green).

**Fig. 8.**
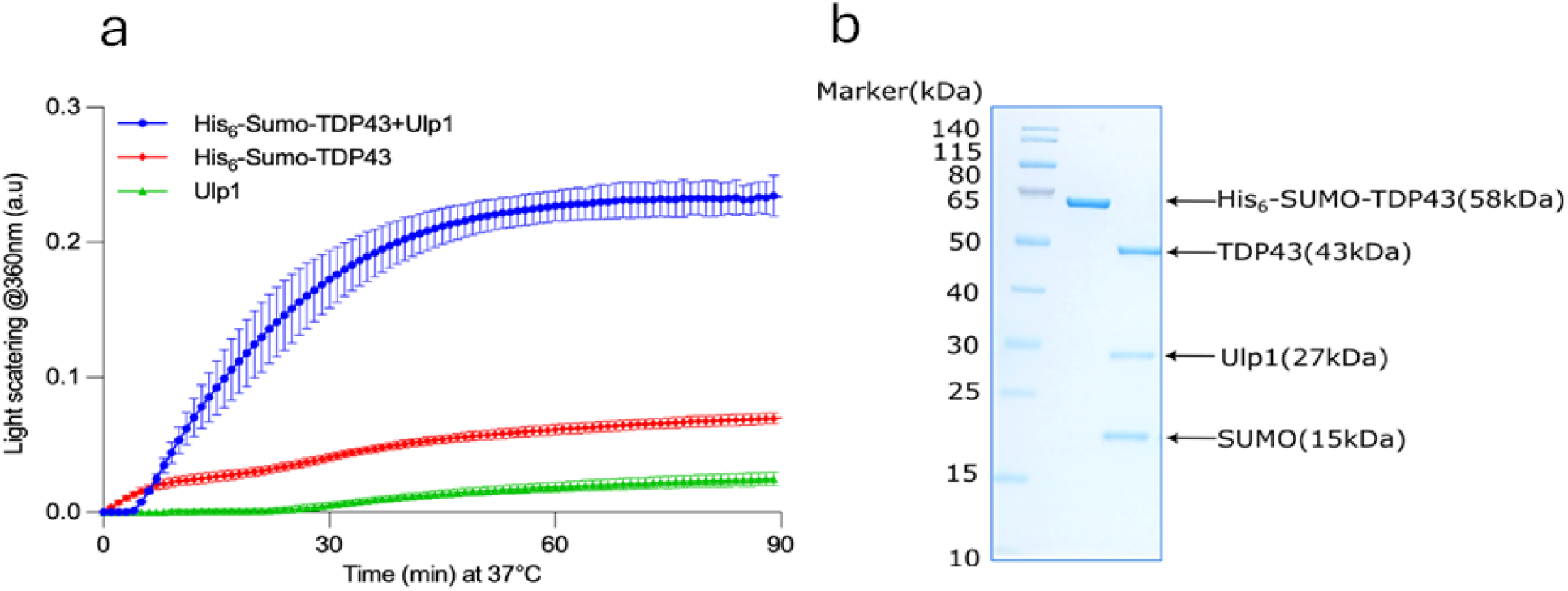
Light scattering analysis of SUMO-cleaved TDP43 aggregation. The aggregation propensity of TDP43 upon SUMO-tag cleavage was assessed using a light scattering assay. The reaction was initiated by adding Ulp1 protease to a 5 µM solution of His_6_-SUMO-TDP43 in a 384-well black, clear-bottom plate. The sample was incubated at 37℃, and light scattering was monitored for 120 minutes at a wavelength of 360 nm in a TECAN Spark plate reader. The data plotted is representative of three biological repeats in GraphPad prism (b) Coomassie-stained 12% SDS-PAGE of the reaction mixture from panel a, showing bands for His₆-SUMO–TDP43 (∼58 kDa), TDP-43 (∼43 kDa), Ulp1 (∼27 kDa), and SUMO (∼15 kDa).

## Discussion and Concluding Remarks

The biochemical study of TDP-43 is essential for understanding its role in ALS/FTD and related proteinopathies and for developing targeted therapeutic strategies. Although several groups have previously purified full-length TDP-43, its recent classification as a prion-like protein now requires work under BSL-2 containment, adding logistical and biosafety constraints. Our protocol is designed to lower these barriers while maintaining rigorous quality control. A key feature is that all size and oligomeric-state information is obtained by single-particle mass photometry rather than SEC-MALS or other fluidic systems (FPLC). Because mass photometry has no internal fluidic path, there is no risk of column or flow-cell carryover by highly aggregation-prone TDP-43, and decontamination is reduced to simple slide and buffer handling within the BSL-2 workflow.

Two additional design choices are critical for the robustness of this purification. First, we intentionally operate the immobilized metal affinity step under resin-limiting conditions to enforce selective capture of His₆-SUMO-TDP43. The soluble fraction contains ∼100 mg total protein, whereas we use only 1.5 ml of Ni-NTA resin with a nominal binding capacity of ∼ 40 mg protein per ml. This ratio promotes competition for binding sites and strongly biases occupancy toward the high-affinity His_6_ tag on SUMO-TDP43, reducing co-purification of weakly interacting host proteins. Second, because our lysis buffer contains 5 mM EDTA and 2 mM DTT, we employ EDTA/DTT-tolerant Ni-NTA resin (Roche, cat. 5893682001), which retains binding capacity in up to 10 mM EDTA, 10 mM DTT, and high concentrations of chaotropes (6 M guanidinium-HCl, 8 M urea; pH 2–14). This compatibility allows us to maintain chelation and reducing conditions throughout lysis and on-column urea refolding without reformulating buffers or risking metal stripping from the resin; conditions under which standard Ni-NTA matrices would fail.

Finally, for buffer exchange and concentration, we deliberately use 100 kDa MWCO centrifugal devices, even though monomeric TDP-43 is ∼55 kDa. Empirically, this MWCO results in minimal protein loss while noticeably reducing aggregation during concentrating, likely by shortening residence time at the membrane and lowering the extent of confinement-induced self-association. Across the workflow, protein integrity (anti-TDP-43 Western blot), oligomeric state (mass photometry), and function (fluorescence anisotropy nucleic-acid binding) consistently meet predefined acceptance criteria for native protein. Together, these features make the protocol safe, reproducible, and technically accessible for routine production of full-length TDP-43 in standard BSL-2 laboratories.

## Safety Procedures for Handling Prion-Like Proteins, in this case : TDP-43

- Containment & Scope: All work with purified, recombinant TDP-43 (Risk: proteopathic seeding) must be performed by trained personnel within a designated BSL-2 laboratory, using double gloves, lab coat, and eye protection.
- Transport & Handling: All movement of samples between BSL-2 and BSL-1 areas must utilize sealed primary tubes within a labeled, rigid secondary container. A transport and waste log must be maintained.
- Liquid Waste Decontamination: All liquid waste containing TDP-43 must be collected and inactivated by mixing with 1 M NaOH (final concentration) for a minimum contact time of > 60 minutes before neutralization and disposal.
- Surface & Spill Decontamination: Solid surfaces and spills must be decontaminated by flooding with 1% Hellmanex III, 2% SDS, or 1 M NaOH for a minimum contact time of > 60 minutes, followed by thorough rinsing or collection as BSL-2 waste.
- Solid Waste Disposal: All contaminated consumables (e.g., columns, beads, and mass photometry (MP) slides) must be collected and discarded as segregated, labeled BSL-2 solid waste in 30L or 60L blue boxes (certified UN 3291). Where practical, items should be soaked in 2% SDS for > 60 minutes before disposal.

## Key Resources Table: Reagents and instruments

| REAGENT or RESOURCE | SOURCE | IDENTIFIER |
| --- | --- | --- |
| Antibodies |  |  |
| Anti-TDP43 polyclonal rabbit antibody | Invitrogen | Cat# PA5-29949 |
| Anti-rabbit HRP-conjugated secondary | Cytiva Amersham | NA934 |
| Bacterial Strains |  |  |
| E. coli BL21-CodonPlus (DE3)-RIPL | Agilent | Cat# 230280 |
| **Chemicals** |  |  |
| Phenylmethylsulfonyl fluoride (PMSF) | Sigma Aldrich | Cat# 93482-50ml-F |
| cOmplete™ Protease Inhibitor Cocktail | Roche | Cat# 11697498001 |
| Lysozyme | Roche | Cat#45-10837059001 |
| DNase I | Roche | Cat# 11284932001 |
| RNase A | Sigma Aldrich | Cat# R4875 |
| EDTA-compatible Ni<sup>2+</sup>-NTA beads | Roche | Cat# 05893801001 |
| ECL Western Blotting Analysis System | Cytiva | Cat# RPN2108 |
| Hellmanex III | Sigma Aldrich | Cat# Z805939 |
| OmniPure water | Sigma Aldrich | Cat# 9601-100ml |
| Bradford Reagent | ThermoFisher | Cat# 1856209 |
| Experimental Models: Plasmids |  |  |
| pET28(a)-His<sub>6</sub>-SUMO-FLTDP43 | GeneScript | This paper |
| **Oligonucleotides** |  |  |
| ssDNA FAM-linked (TG)<sub>4</sub> | Sigma | Custom |
| ssDNA FAM-linked (AC)<sub>4</sub> | Sigma | Custom |
| **Instruments** |  |  |
| Microfluidizer | LM20 |  |
| iBRIGHTFL1000 | Invitrogen |  |
| TwoMP Mass Photometer | Refeyn |  |
| Tecan Spark plate reader | Tecan |  |
| ChirascanV100 | Applied photophysics |  |
| **Other** |  |  |
| 0.45μm PVDF filter | Merck | Cat# SLHV033NS |
| 0.2 μm PES filter | Merck | Cat# SCGP00525 |
| 2ml protein low binding tubes | Eppendorf | Cat# 022431102 |
| Amicon filter (100 kDa MWCO) | Millipore/Merck | Cat# UFC910024 |
| 1.5ml Protein LoBind tube | Eppendorf | Cat# 022431081 |
| 12% mPAGE BIS-TRIS precast gels | Merck | Cat# MP12W12 |
| 4x LDS sample buffer | Merck | Cat# MSPSB-60ml |
| Pre-stained protein ladder (10-140 kDa) | ThermoFisher | Cat# 26616 |
| mPAGE™ MES SDS running buffer | Merck | Cat# MPMES |
| QuickBlue protein stain | LuBioScience GmbH | Cat# LU001000 |
| Nitrocellulose membrane (0.4 μm) | Bio-Rad | Cat# 1620145 |
| 10x-Tris Glycine Transfer buffer RunBlue™ | Abcam | Cat# ab270518 |
| MassGlass™ UC sample prep kit | Refeyn | Cat# MP-CON-21022 |
| 0.2 μm syringe filter | Merck | Cat# SLGVX13NL |
| 1 mm path-length quartz cuvette | Hellma | Cat# 110-1-40 |
| 384-well black clear-bottom plate | Greiner | Cat# 781906 |
| Optical adhesive covers | ThermoFisher | Cat# 4360954 |

## Acknowledgment

We would like to express our gratitude to the members of Prof. Henning Stahlberg’s lab, particularly Dr Babatunde Ekundayo and Kazedi Ekundayo, for their invaluable training and essential support in conducting experiments within the BSL-2 facility and Prof. Sahand Jamal Rahi’s (LPBS) lab to allow us to use lab instruments. S.T. acknowledges SNSF funding under grant CRSII5_193740.

The authors declare no conflicting interests.

## Contributions

S.T. and P.D.L.R. conceived the project. S.D, S.T., and P.D.L.R. designed the experiments. S.D, and S.T. generated reagents (plasmids and proteins). S.D., and S.T performed the experiments. S.D., S.T., and P.D.L.R. analysed the data and created the figures. S.D, S.T., and P.D.L.R. wrote the manuscript. All authors read, edited, and approved the manuscript.

## Notes

### Competing Interest Statement

The authors have declared no competing interest.

